# Selective Corticofugal Modulation on Sound Processing in Auditory Thalamus of Awake Marmosets

**DOI:** 10.1101/2021.03.13.435231

**Authors:** Yuanqing Zhang, Xiaohui Wang, Lin Zhu, Siyi Bai, Rui Li, Hao Sun, Runze Qi, Ruolan Cai, Min Li, Guoqiang Jia, Kenneth E Schriver, Xinjian Li, Lixia Gao

## Abstract

Cortical feedback has long been considered crucial for modulation of sensory processing. In the mammalian auditory system, studies have suggested that corticofugal feedback can have excitatory, inhibitory, or both effects on the response of subcortical neurons, leading to controversies regarding the role of corticothalamic influence. This has been further complicated by studies conducted under different brain states. In the current study, we used cryo-inactivation in the primary auditory cortex (A1) to examine the role of corticothalamic feedback on medial geniculate body (MGB) neurons in awake marmosets. The primary effects of A1 inactivation were a frequency-specific decrease in the auditory response of MGB neurons coupled with an increased spontaneous firing rate, which together resulted in a decrease in the signal-to-noise ratio. In addition, we report for the first-time that A1 robustly modulated the long-lasting sustained response of MGB neurons which changed the frequency tuning after A1 inactivation, e.g., neurons with sharp tuning increased tuning bandwidth whereas those with broad tuning decreased tuning bandwidth. Taken together, our results demonstrate that corticothalamic modulation in awake marmosets serves to enhance sensory processing in a way similar to center-surround models proposed in visual and somatosensory systems, a finding which supports common principles of corticothalamic processing across sensory systems.

## Introduction

For survival, it is vital for mammals to detect ambient sounds rapidly and accurately. This is largely accomplished by interaction of the ascending and descending primary auditory pathways (Nunez and Malmierca, 2007). In order to detect sounds in a complex acoustic environment, previous studies have indicated the importance of the descending corticothalamic pathway in the control of auditory information flow (Crick, 1984; Harth et al., 1987; He, 1997; Murphy and Sillito, 1987; Sillito et al., 1994; Villa et al., 1991), something which we hypothesize may be mediated by enhancing driven response of auditory neurons as well as by suppressing spontaneous activity. The corticothalamic projection is about 10 times more extensive than the thalamocortical projection (Andersen et al., 1980; Guillery, 1967; Liu et al., 1995; Montero, 1991; Winer et al., 2001; Winer and Larue, 1987), which has been suggested to serve as a gating or gain control in the processing of sensory information from the periphery to the cortex (Crick, 1984; Harth et al., 1987; He, 1997; Murphy and Sillito, 1987; Sillito et al., 1994; Villa et al., 1991). Several studies of visual and barrel somatosensory systems proved that the corticothalamic projection may increase the sensory driven response in thalamic neurons by enhancing point-to-point homologous projections and suppressing non-homologous projections (Kalil and Chase, 1970; Macchi et al., 1986; Singer, 1977; Temereanca and Simons, 2004; Tsumoto et al., 1978; Usrey and Sherman, 2019; Yuan et al., 1985, 1986). However, whether and how corticothalamic projections modulate auditory processing is far less clear (Nunez and Malmierca, 2007; Terreros and Delano, 2015; Wang, 2007). While previous studies have used focal activation and inactivation methods to study the effect of corticothalamic feedback, there have been few studies examining modulation of cortical feedback at a global scale in awake animals, something which may be particularly crucial for establishing spontaneous activity levels in dynamic environments.

There is accumulating evidence showing that a remarkable feature of A1 neurons in awake animals is the long-lasting sustained response to preferred stimuli (Gao and Wang, 2019; Wang et al., 2005), distinct from the transient responses observed in anesthetized animals (Heil, 1997; Phillips, 1985). However, most of the previous corticofugal studies in auditory system were carried out in anesthetized animals (Amato et al., 1969; He, 1997, 2003b; He et al., 2002; Ryugo and Weinberger, 1976; Villa et al., 1991; Watanabe et al., 1966), where the auditory cortex was suppressed to some degree (He, 1997). In this case, manipulation of A1 demonstrated some degree of modulation of spectral, intensity and spatial aspects of onset response during auditory processing (He, 2003a; Nunez and Malmierca, 2007; Suga, 2020; Terreros and Delano, 2015; Winer, 2006; Winer et al., 2001). Although sustained responses are crucial for temporal processing (Wang et al., 2008), and especially important for speech and music processing (Rosen, 1992; Wang, 2000), the effect of corticothalamic modulation on sustained response remains unknown. Due to the relatively delayed effect of corticofugal modulation (in comparison to direct sensory response), the effects of corticothalamic modulation may be especially evident on the long-lasting sustained response, raising the possibility that there are temporal effects that have not yet been examined.

In this study, to examine the effect of corticothalamic feedback on spontaneous firing rate and evoked response as well as on the sustained response in MGB neurons, we inactivated A1 of awake marmosets via cryo-inactivation. By examining how A1 modulates the spectral and temporal response of MGB neurons, we found inactivation of A1 decreased not only the auditory response but also the SNR of MGB neurons. Moreover, our results showed that effects of A1 inactivation were more prominent on the sustained response than on the onset responses of MGB neurons, which led to changes in the frequency tuning bandwidth of MGB neurons. The implications of these results are discussed.

## Results

### Validation of Cortical Inactivation via Cryoloop

To investigate whether and how corticothalamic projection affects sound processing of MGB neurons, we inactivated A1 of awake marmosets reversibly and chronically via a cryoloop (Fig. 1C), based on previous methods (Chen and Stuphorn, 2018; Chen et al., 2020; Lomber and Malhotra, 2008; Lomber et al., 1999; Petersen and Buzsaki, 2020; Ryugo and Weinberger, 1976; Villa et al., 1991; Yu et al., 2016). Although the cryoloop has been extensively used to inactivate neural activity of various brain regions in rodents (Petersen and Buzsaki, 2020), cats (Lomber and Malhotra, 2008; Lomber et al., 1999; Ryugo and Weinberger, 1976; Villa et al., 1991; Yu et al., 2016) and other primates (Chen and Stuphorn, 2018; Chen et al., 2020), it has never been tested in awake marmosets. To test the effectiveness of our cooling system, a cryoloop with a temperature sensor was placed on the surface of A1 of anesthetized rats. We found sound elicited spiking responses vanished when the temperature fell to 3 ℃ and were restored when the temperature was raised back to a normal temperature (Fig. S1A-B). The inactivated brain area was within 2.5 × 2.5 mm in distance and depth at a temperature of 3 °C (Fig. S1C-J).

**Figure 1.**
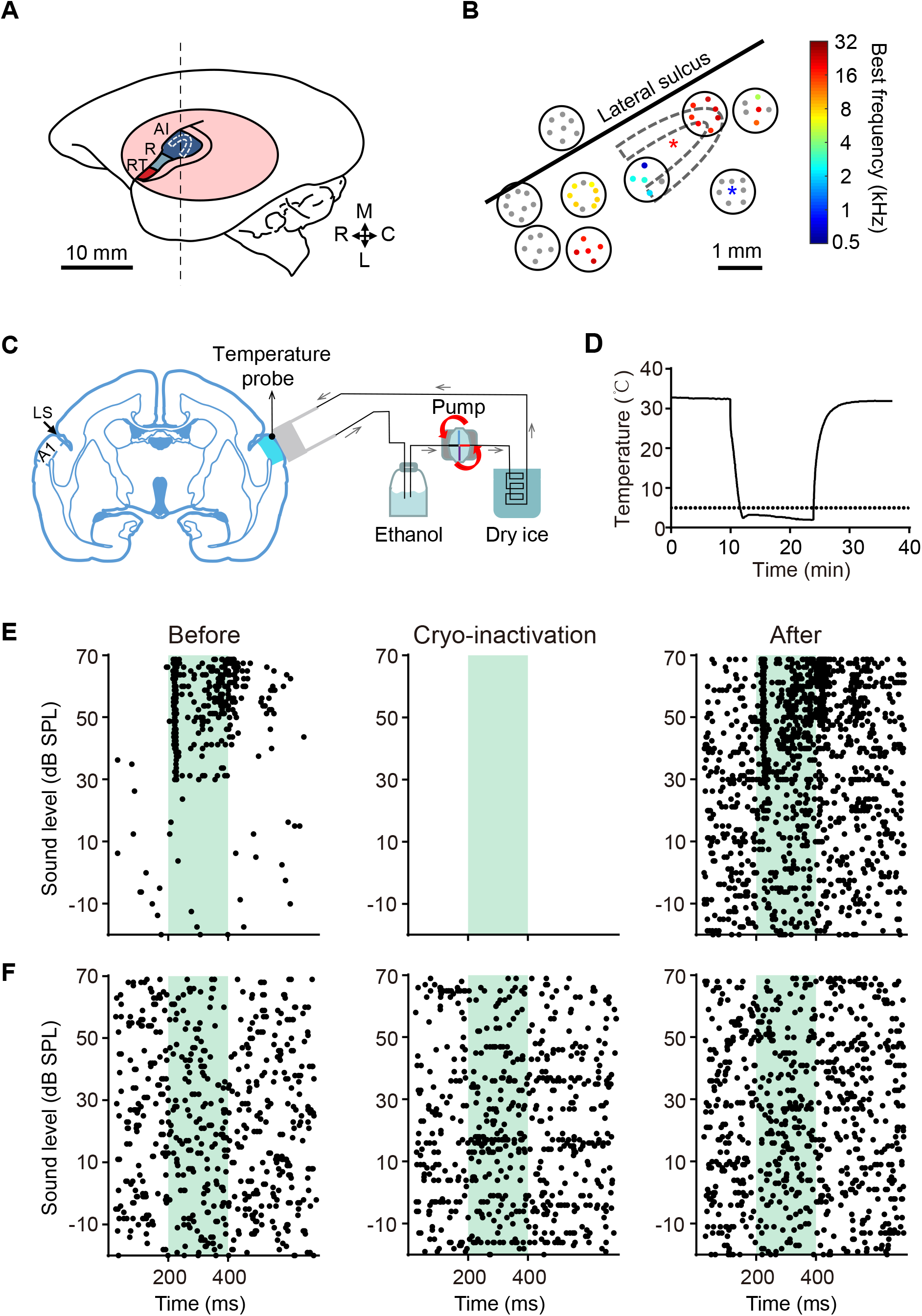
Validation of Cooling Method for Inactivation of A1 in Awake Marmosets. (**A**) Sketch map showing the location of auditory cortex and relative position of cryoloop (U-shaped white dotted loop) in left hemisphere of marmoset monkeys. The core regions of auditory cortex are presented by different colors, which includes the primary auditory cortex (A1) and rostral fields (R and RT). The pink oval indicates the position of the recording chamber. The dashed vertical line shows the position for the coronal section in C. R, rostral; C, caudal; M, medial; L, lateral. Scale bar: 10 mm. (**B**) An example showing best-frequency map in the auditory cortex (animal ID: BJ1612F1, left hemisphere) and the relative position of cryoloop in the auditory cortex. The black circles indicate the position of miniature holes (diameter, 1mm) through which the cortical neurons were accessed. Pure-tone-responsive recording sites are indicated by colored dots showing the spatial distribution of best frequency (BF). Pure-tone-nonresponsive recording sites are indicated by grey dots. The location of A1 was identified by the tonotopic gradient (Rostral: low frequency; Caudal: High frequency) and reversal of BF on the boundary. The asterisks show the penetrations through which single unit recordings were performed to validate the cooling method in E and F. Scale bar: 1 mm. (**C**) Schematic diagram of the cooling device. The black dot shows the position of the temperature probe. The flow speed can be adjusted by the pump. The arrows indicate direction of fluid flow. LS, lateral sulcus. (**D**) Temperature change detected by the temperature probe in an experiment. The dashed line indicates 5 °C. (**E-F**) Raster plots showing the neural spiking responses to white broad-band noise (WB) with different sound levels before, during and after inactivation of A1. The recording sites are indicated by the asterisks in B (E, Red; F, Blue). Green shaded areas indicate periods of acoustic stimulation.

To further validate the inactivation effect, a stainless-steel cryoloop was positioned on the surface of A1 of awake marmosets after the location of A1 was identified by its response characteristics and tonotopic gradient (Fig. 1A-B). Meanwhile, a temperature sensor touched with the brain surface was used to detect temperature changes in A1 (Fig. 1C-D). Similar to the results in anesthetized rats, spiking activity adjacent to the cryoloop of awake marmosets was totally blocked when the temperature fell to 3 °C and restored at normal temperature (Fig. 1 E). If the recording location was 1 mm or more away from the cryoloop, the neural spiking activity was not affected by A1 cooling (Fig. 1F). After validation of the inactivating effect, the cryoloop assembly was implanted into the recording chamber chronically. It is important to note that the distance between MGB and the implanted cryoloop was > 8 mm. As a result, there was no change in spiking activity in > 50% of MGB neurons during inactivation of A1. This suggests a lack of direct inactivation of MGB by the cryoloop (Fig. 2H).

**Figure 2.**
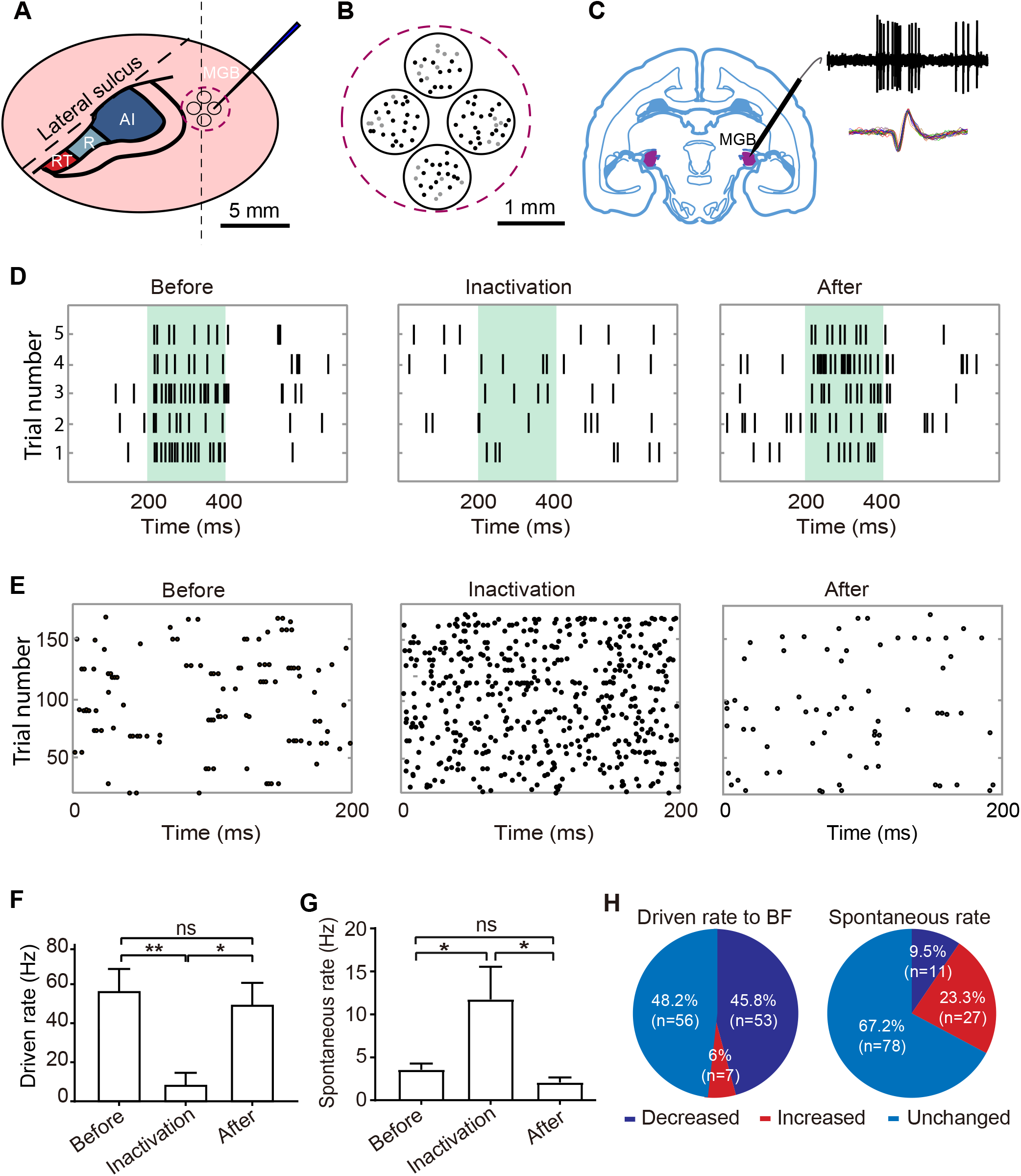
Neural Activity of MGB Neurons When A1 Was Inactivated. (**A**) Diagram showing relative position for single unit recordings to access MGB neurons. Scale bar: 5 mm. (**B**) Diagram showing the penetrations with auditory (Black) and non-auditory response (Grey) in MGB. Scale bar: 1 mm. (**C**) Location of MGB (purple) in a coronal section made at the dashed line location in A and trajectory of single unit recordings. Subpanels on the top right shows a sampled raw trace recorded by single unit recording (Top) and spike waveforms of this unit (Bottom). (**D**) Raster plots of response to BF tone in a sampled MGB neuron before, during and after inactivation of A1. Green shaded areas indicate periods of acoustic stimulation. (**E**) Raster plots of spontaneous activity in a sampled MGB neuron before, during and after inactivation of A1. (**F-G**) Driven rate in D and spontaneous firing rate in E before during and after inactivation of A1. * indicates p<0.05, ** indicates p<0.01, ns indicates no significant difference. (**H**) Proportion of MGB neurons with increased, decreased, and unchanged sound driven response to BF tones (Left) as well as increased, decreased, and unchanged spontaneous rate when A1 was inactivated, respectively.

### Inactivation of A1 Mainly Decreased Auditory Response of MGB Neurons and Increased Their Spontaneous Activity

Based on a previous report (Bartlett and Wang, 2011) and guided by structural Magnetic Resonance Imaging (MRI), a tungsten electrode was used to access MGB at an angle of 57.5° and 3-mm lateral to the high-frequency representation of A1 via 1mm-diameter craniotomy (Fig. 2A-C). Out of 278 recorded neurons, 116 neurons fulfilled the criteria of being well-isolated, recorded before, during and after inactivation of A1, and exhibited recovery of spiking activity after cooling; this population was used for further data analysis. In the auditory system, spontaneous activity levels representing background noise is particularly important for sound processing. To assess the effect of corticofugal modulation, both sound-driven and spontaneous activity were analyzed. We found inactivation of A1 decreased sound elicited responses of MGB neurons (Fig 2D, 2F; P=0.008) and increased spontaneous firing (Fig 2E and 2G; P =0.025). 45.8% (53/116) of MGB neurons exhibited a significant decline in the sound driven response, and 6% (7/116) exhibited an increase in sound driven response (Fig 2H, Left). This revealed that under awake conditions A1 has a primarily facilitatory effect on the response of MGB neurons to auditory stimuli. The result is consistent with that in the auditory system of bats (Zhang et al., 1997) as well as in the visual system and the somatosensory barrel system (Kalil and Chase, 1970; Macchi et al., 1986; Singer, 1977; Tsumoto et al., 1978; Yuan et al., 1985, 1986). Interestingly, with respect to spontaneous firing rate, 23.3% (27/116) of MGB neurons exhibited an increase and 9.5% (11/116) exhibited a decrease (Fig 2H, Right). These results indicated that in awake state A1 modulates both sound evoked response and spontaneous activity, and that these effects are distinct.

### Inactivation of A1 decreased the signal-to-noise ratio (SNR) of MGB neurons

The results above suggest that A1 modulates auditory response and spontaneous activity of MGB neurons in different ways. We then hypothesized that this differential effect would lead to increase of the SNR of MGB neurons. To test this, we analyzed spontaneous firing rate and sound driven response of individual MGB neurons in response to tone of varying frequency. We found that the frequency tuning of MGB neurons was broadened during inactivation of A1 as shown by the example neuron (Fig.3A-B). This broadening consisted of two components. During inactivation of A1, the driven response of this neuron largely decreased when presented tones across its preferred frequencies, especially to its best frequency (BF, Fig. 3B-C). In contrast, its spontaneous firing increased across a broad range of frequencies (Fig. 3B, D). Overall, inactivation of A1 resulted in opposing effects on sound driven response vs spontaneous firing rates (Fig 3E-F).

**Figure 3.**
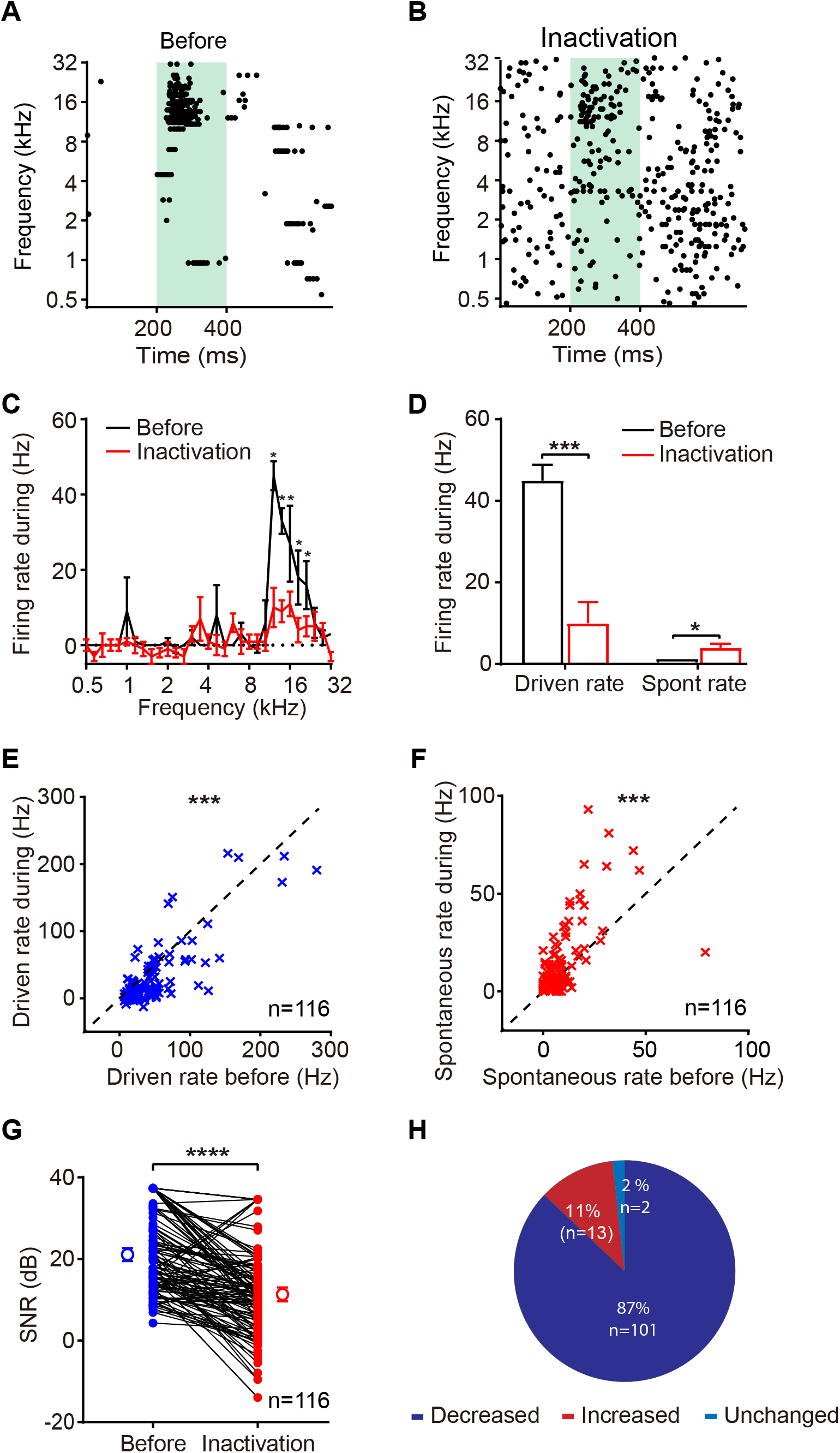
Inactivation of A1 Decreased the Signal-to-Noise Ratio of MGB Neurons. (**A-B**) Raster plots showing the neural response of an example MGB neuron to pure tones with varying frequencies before (A) and during (B) inactivation of A1. Green shaded areas indicate periods of acoustic stimulation. (**C**) Frequency tuning curve of neuron in A-B before and during inactivation of A1. Analysis window is period of acoustic stimulation (200-400 ms). (**D**) Driven rate and spontaneous rate (spont rate) of neuron in A-B before and during inactivation of A1, respectively. * indicates p<0.05, *** indicates p<0.001. (**E-F**). Change of driven rate (E) and spontaneous rate (F) of all MGB neurons before and during inactivation of A1, the dashed lines are diagonal lines. *** indicates p<0.001. (**G**) Scatter plot showing signal-to-noise ratio (SNR) of MGB neurons before and during inactivation of A1. **** indicates p<0.0001. (**H**) Proportion of MGB neurons with increased, decreased, and unchanged SNR caused by inactivation of A1. Change of ±0.05 dB was considered a significant difference.

Next, we calculated the SNR (SNR=20 log_10_ (driven rate/spontaneous rate), see Methods) of 116 MGB neurons before and during inactivation of A1. 87% (101/116) of MGB neurons exhibited a significant decrease in SNR after inactivation of A1(Fig 3G-H). These results demonstrated that with the sound stimuli presented, in awake marmosets, the primary modulatory effect of corticothalamic feedback is an increase in the SNR of MGB neurons, something which may be beneficial to the detection of target sounds.

### Stimulus-dependent Corticofugal Modulation of MGB neurons

Given that corticofugal modulation increased SNR of MGB neurons during sound processing, we wondered whether the corticofugal modulation exhibits specificity for the features of acoustic stimuli (e.g. spectral, intensity and temporal information). To examine this, we presented acoustic stimuli with varying frequency, intensity, and modulation frequency (MF) to drive MGB neurons before, during and after inactivation of A1. We found that, in contrast to non-BF tones which elicit transient onset responses, the example MGB neuron (Fig. 4A) exhibited onset response followed by a long-lasting sustained response to BF tones (around 16 kHz, in red box). Interestingly, inactivation of A1 resulted mostly in a decrease of neural activity to BF tone (Fig. 4D), without change of temporal firing pattern for non-BF tones (Fig. 4A). For the same MGB neuron, inactivation of A1 caused no change of temporal firing pattern either to intensity-varying (Fig. 4B) or to time-varying stimuli (sinusoidal amplitude modulated tones, Fig. 4C). These results indicated that the effect of corticofugal modulation varied with the features of acoustic stimulation (Fig. 4D). To further confirm these results, we examined the specificity of stimulus effect in 24 of the 116 MGB neurons (Fig.4E). We found inactivation of A1 led to varying degrees of MGB modulation to sound features (Fig. 4E-F). Among them, 13 of 24 (54%) neurons showed either facilitatory or suppressive effect to varying acoustic stimuli, and 46% of neurons showed only facilitation or suppression to different acoustic stimuli (Fig. 4G). Thus, with the stimuli tested, roughly half of the neurons exhibited feature specific effects. We speculate that, if more stimuli with other features were tested, additional varying effects of modulation may be induced. Taken together, our results indicate that corticofugal modulation on MGB neurons is stimulus-dependent.

**Figure 4.**
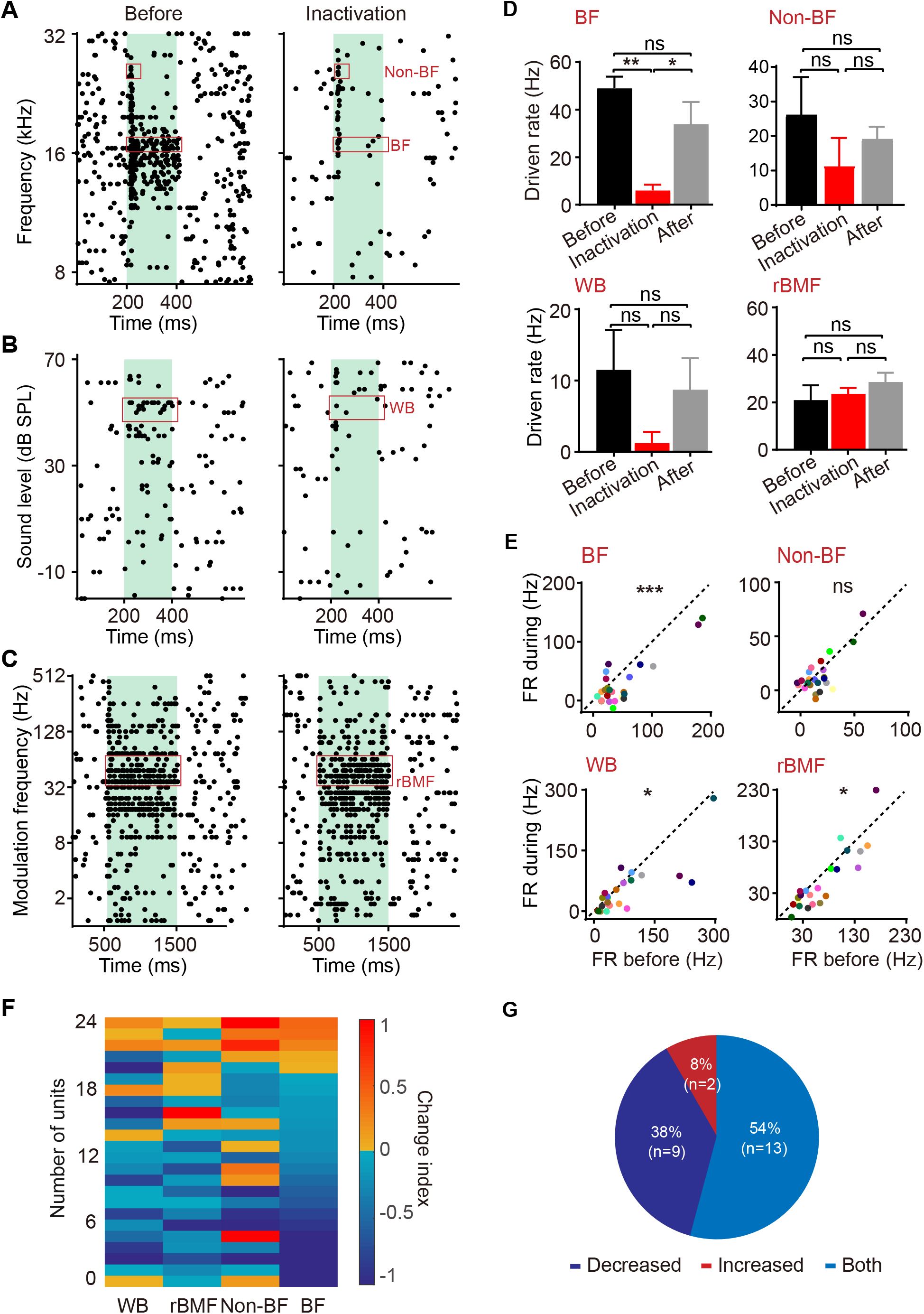
A1 Modulates Evoked Response of MGB Neurons in a Stimulus-Dependent Manner. (**A-C**) Raster plots of an example MGB neuron in response to pure tones with varying frequency (A), white broad-band noise (WB) with varying intensity (B) and an amplitude modulated tone (sAM) with varying modulation frequency (C) before and during inactivation of A1. The green shaded region indicates periods of acoustic stimulation. The red boxes represent analysis window in D. BF: Best Frequency; Non-BF: the frequency which elicited the smallest statistically significant driven response; rBMF: modulation frequency of sAM which elicited the maximal driven response. (**D**) Statistics analysis showing driven rate of the neuron in A-C in response to BF, Non-BF, WB and rBMF before, during and after inactivation of A1. The analyses windows were indicated by the red box in A-C. * indicates p<0.05, ** indicates p<0.01, ns indicates no significant difference. (**E**) Scatter plots showing the driven rate of 24 MGB neurons in response to BF, Non-BF, WB and rBMF before and during inactivation of A1, respectively. X-axis: firing rate before inactivation; Y-axis: firing rate during inactivation. The dashed lines indicate complete correlation. (**F**) Pseudo-color image showing the changes of driven rate of MGB neurons to different stimuli. The color indicates the value of change index (see Methods). (**G**) Proportion of MGB neurons showing increased, decreased or both increased and decreased responses to different sound stimuli when A1 was inactivated.

### Nonlinear Corticofugal Modulation on Spectral Processing of MGB neurons

We then hypothesized that, due to the time lags inherent in thalamo-cortico-thalamic (MGB to A1 to MGB) loops, corticothalamic modulation may be differentiated by the onset and by the long-lasting response of MGB neurons in awake animals. To test this hypothesis, the corticothalamic modulation effect was measured in MGB neurons with long-lasting sustained response to BF tone. Strikingly, inactivation of A1 changed the temporal responses of MGB neurons around BF tones, revealing a switch from either a long-lasting sustained response to a transient onset response (Fig.5A) or vice versa (Fig. 5B); this was especially prominent at the BF tone. As a result, firing rate and temporal pattern following presentation of BF tones significantly changed during inactivation of A1(Fig. 5C-D, Left).

**Figure 5.**
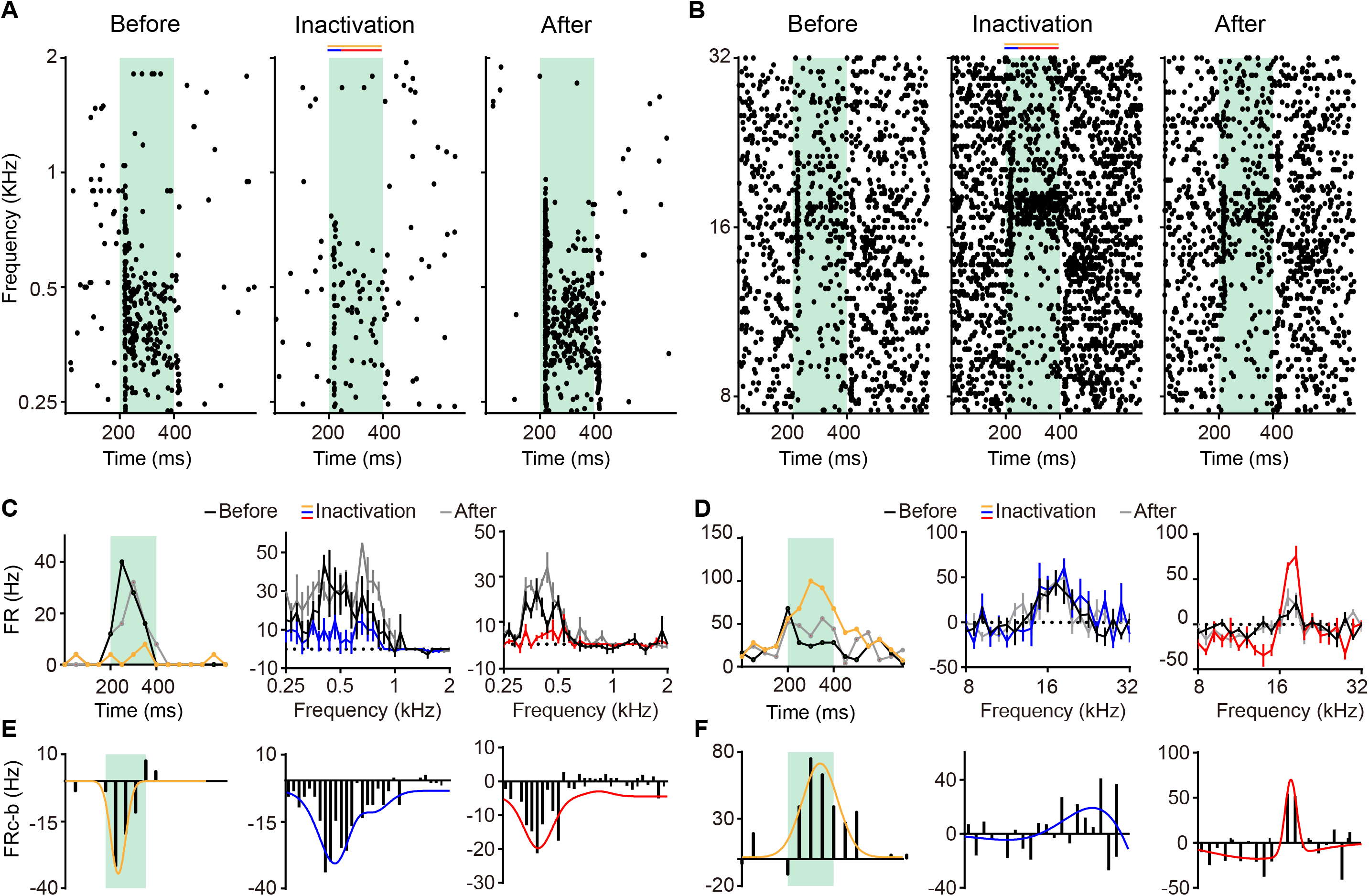
Inactivation of A1 Modulated the Frequency Tuning of MGB Neurons. (**A-B**) Raster plots of two example MGB neurons in response to pure tones with varying frequencies before (Left), during (Middle) and after (Right) inactivation of A1. Green shaded areas indicate periods of acoustic stimulation. The color bars at the top indicate the analyses window of onset (200-240 ms, blue), sustained response (240-400 ms, red) or entire stimulus window (200-400 ms, orange). (**C-D**) Left panels show the peri-stimulus time histograms (PSTHs) of the neurons in A and B in response to BF tones before, during and after inactivation of A1. Middle (analyses window: 200-240 ms) and Right (analyses window: 240-400 ms) panels show frequency tuning curves of the neurons in A and B before, during and after inactivation of A1, respectively. **(E-F)** Firing rate (FR) difference during and before A1 inactivation conditions corresponding to figures in C and D above. The lines are Gaussian curve fitting. The color bars on the top applied to C-F, which indicate the analyses windows that are the same as A-B. The line plots indicate the results in corresponding color-coded analyses windows.

To further analyze whether A1 modulates the onset and sustained responses of MGB neurons differently, the frequency tuning curves of MGB neurons were plotted using the onset response (0-40 ms from the stimulus onset, Fig 5C-D, Middle) and sustained response (40-200 ms from the stimulus onset, Fig. 5C-D, Right), respectively. For the same MGB neurons, inactivation of A1 caused more dramatic change in the frequency tuning curves calculated by sustained response than those calculated by onset response (Fig. 5C-D, Middle and Right). The modulation around BF is larger than for frequencies far away from BF which demonstrated that the effect on frequency tuning is not simply a broad gain effect (Fig 5E-F). Notably, the frequency band modulated by A1 was narrower in the late phase sustained response than that in early phase onset responses (Fig 5E-F, Middle and Right). In a few cases (10%, 12/116), inactivation of A1 resulted in loss of frequency tuning of MGB neurons, as indicated by a loss of tuning properties (Fig. S2). These data suggest that A1 exerts not a simple gain modulation on MGB neurons across the frequency spectrum.

### Quantification of Corticofugal Modulation on Temporal and Spectral features

To quantify corticofugal modulation on temporal and spectral features, we compared the sound driven rate of both onset responses (0-40 ms from the stimulus onset, Fig 6A) and sustained response (40-200 ms from the stimulus onset, Fig 6B), before and during inactivation of A1. Both onset and sustained responses decreased firing rate when A1 was inactivated. Next, we calculated the change index of onset response and sustained response, respectively (Fig. 6C). We found inactivation of A1 caused a greater change of sustained response than that of onset response in 48% of MGB neurons; and a smaller change of sustained response than that of onset response in 37% of neurons (Fig. 6D). Next, we compared the tuning bandwidth before and during inactivation of A1 (Fig. 6E). Except for 13 neurons (black circles) which either lost (Fig. S2, n=12) or gained (n=1) frequency tuning during inactivation of A1, a significant fraction of MGB neurons (48.5%) exhibited a tuning bandwidth change of more than 0.2 octaves (decrease: 23%; increase: 25.2%; Fig. 6I, Left). Interestingly, for MGB neurons with changed tuning bandwidth, inactivation of A1 increased the tuning bandwidth of MGB neurons with sharp tuning (0.77 ± 0.10 octaves) and decreased the tuning bandwidth of MGB neurons with broad tuning (1.62 ± 0.23 octaves, Fig 6G). However, whether and how the tuning bandwidth of MGB neurons was modulated by A1 was not predicted by their BF (Fig. 6H). A proportion (31%) of MGB neurons exhibited a BF shift (Fig 6F, 6I, Right), consistent with previous studies (Villa et al., 1991; Zhang et al., 1997). These results demonstrate that A1 feedback modulation sharpened frequency tuning of MGB neurons with sharper tuning and broadened those with broader tuning. This suggests corticothalamic modulation may serve to increase either the frequency sensitivity or frequency response range of MGB neurons.

**Figure 6.**
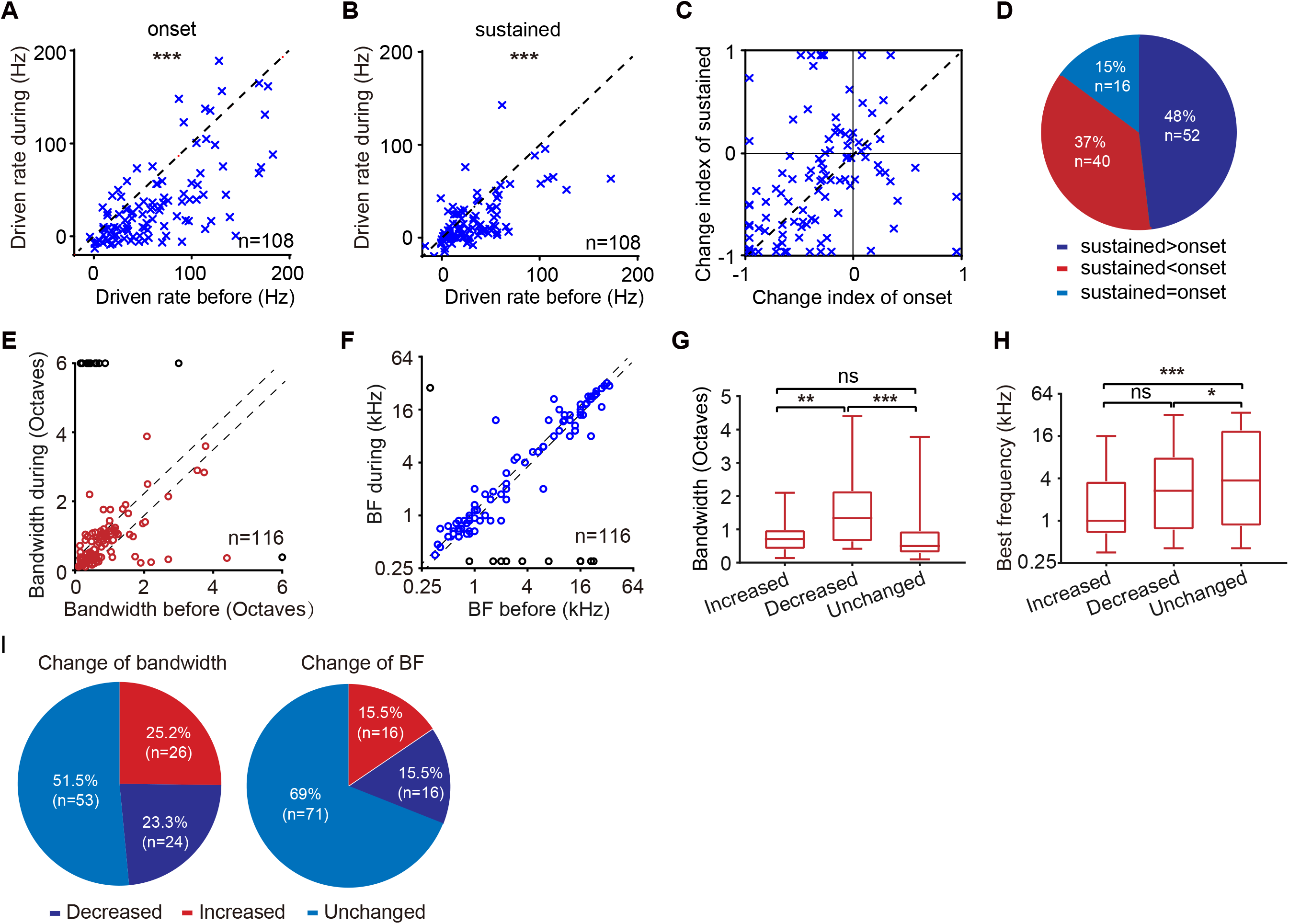
Quantitative Analysis of Temporal and Spectral Change of MGB Neurons Caused by A1 Inactivation. **(A-B)** Driven rate of onset (A) and sustained response (B) following inactivation of A1 plotted against driven rate before inactivation of A1. *** indicates p<0.001. (**C**) Change index of sustained response plotted against that of onset response for 108 MGB neurons. (**D**) Proportion of MGB neurons with change index of sustained response larger, smaller, and equal to that of onset response caused by inactivation of A1. (**E**) Half tuning bandwidths of MGB neurons before inactivation of A1 plotted against those during inactivation of A1. The analyses window is the period of acoustic stimulation (200-400 ms). For MGB neurons showing similar response to different tones and without clear half-bandwidths in some conditions, a value of 6 octaves was assigned and indicated by the black circles. The dashed lines indicate the confidence interval (0.2 octaves). (**F**) BF shift of MGB neurons during inactivation of A1. For MGB neurons without a clearly identified best frequency, a fixed value (0.3 kHz) is assigned as BF and indicated by the black circles. The dashed lines indicate the confidence interval (0.2 octaves). (**G**) Average tuning bandwidth of MGB neurons with decreased, increased, and unchanged tuning bandwidth caused by A1 inactivation, respectively. (**H**) Average BFs of these three groups of MGB neurons as shown in E. (**I**) Proportion of MGB neurons exhibiting increased, decreased, and unchanged tuning bandwidth (Left) or BF (Right) as shown in E and F, respectively.

## Discussion

The importance of corticofugal modulation on auditory processing and perception is well recognized (He, 2003b; Nunez and Malmierca, 2007; Suga, 2020; Terreros and Delano, 2015; Winer, 2006; Winer et al., 2001; Wood et al., 2017). However, previous studies which applied cortical activation or inactivation have found that corticofugal influence can be either suppressive or facilitatory, leading to controversy in the field (He, 2003b; Suga, 2020). Here, we hypothesized that some of these apparently inconsistent results are due to the fact that these studies were largely conducted in anesthetized animals (Aitkin and Dunlop, 1969; Amato et al., 1969; He, 2003a; Ryugo and Weinberger, 1976; Villa et al., 1991; Watanabe et al., 1966). In the present study, following reversible inactivation of A1 via cryo-inactivation in *awake* marmosets, we found inactivation of A1 led to an increase in spontaneous firing rate and a decrease in the driven rate of MGB neurons (Fig. 2 and Fig. 3). Furthermore, we found that the sustained response of MGB neurons at or near their BF vanished or appeared during A1 inactivation, indicating an influence on frequency tuning beyond a simple gain modulation (Fig.5 and Fig. 6). Thus, we report two previously unknown findings in the auditory thalamocortical system. We find that first, in the awake animal, corticothalamic feedback plays a positive modulatory role which serves to improve the SNR in auditory processing and that second, corticothalamic feedback shapes the temporal firing pattern in response to auditory stimuli (sustained/transient), something which may be particularly critical for auditory processing in complex sound environments. Importantly, our study in awake animals parallels the profile of thalamocortical circuits proposed in other sensory systems, thereby supporting common principles of thalamocortical function.

### Controversy due to animal’s brain state

Although the role of corticothalamic modulation has been extensively studied, the underlying neural mechanism remains unclear. Most previous studies which were conducted in anesthetized animals, found that activation of auditory cortex decreased the auditory response of MGB neurons (Amato et al., 1969; Watanabe et al., 1966). This suggested an inhibitory role of auditory corticofugal modulation, which serves to limit excitation in a reverberatory thalamocortical loop. In contrast, in the visual system (Andolina et al., 2007; Kalil and Chase, 1970; Macchi et al., 1986; Sininger et al., 1999; Tsumoto et al., 1978; Wang et al., 2018; Yuan et al., 1985, 1986) and in barrel somatosensory system(Ghazanfar et al., 2001; Temereanca and Simons, 2004), studies report an enhancement of point to point homologous projections along with a suppression at non-homologous projections (Kalil and Chase, 1970; Macchi et al., 1986; Singer, 1977; Temereanca and Simons, 2004; Tsumoto et al., 1978; Usrey and Sherman, 2019; Yuan et al., 1985, 1986). This finding has been further supported in a highly specialized auditory system, that of awake bats, where activation of A1 demonstrated improved sound processing at subcortical levels (Suga, 2020). This raises the possibility that differences in previous studies arise from different brain states (Bartlett and Wang, 2007; Constantinople and Bruno, 2011; Haider et al., 2013; Long and Lee, 2012). That is, sound processing in auditory cortex of awake animals may differ fundamentally from that of anesthetized animals (Gao and Wang, 2019; Orman and Humphrey, 1981; Wang et al., 2005), especially in the temporal domain (Wang et al., 2008). While there may be species differences, we propose that the previous contradictory results may be due to brain state-related effects, including depression of responses and effects on temporal aspects of auditory responses by anesthesia (Wang et al., 2008). More specifically, we found corticothalamic modulations decreased spontaneous background activity and increased stimulus driven response, resulting in increased SNR of MGB neurons during sensory processing. Such effects may not be evident in anesthetized studies due the suppressed level of spontaneous activity. In summary, our result is consistent with the auditory studies in bats (Zhang et al., 1997) as well as with the bulk of corticofugal studies in other sensory systems (Andolina et al., 2007; Ghazanfar et al., 2001; Kalil and Chase, 1970; Macchi et al., 1986; Sininger et al., 1999; Temereanca and Simons, 2004; Tsumoto et al., 1978; Wang et al., 2018; Yuan et al., 1985, 1986).

### Stronger Modulation to Long-lasting Sustained Response

Temporal information embedded in sounds is crucial for perceiving and discriminating communication sounds, such as that in human speech (Rosen, 1992) and animal vocalizations (Singh and Theunissen, 2003). In sharp contrast to the transient onset responses which dominate recordings in auditory cortex of anesthetized animals, auditory cortex of awake animals can generate unique sustained firing over the stimulus duration, useful for encoding the temporal information in sound stimuli (Wang et al., 2008). Although it is believed to be important, whether the long-lasting sustained response in MGB is modulated by A1 is still unknown. Due to the time lags inherent in thalamo-cortico-thalamic loops, we therefore hypothesized that, corticothalamic modulation may be differentiated by the onset and by the long-lasting response of MGB neurons in awake animals. We found that effects of A1 inactivation were more prominent on the sustained than on the onset responses of MGB neurons. In addition, long-lasting sustained response of some MGB neurons diminished (Fig. 5A) or emerged (Fig. 5B) when A1 was inactivated. These results confirmed the hypothesis that A1 modulate the onset and sustained response of MGB neurons separately.

### Two Roles of Corticofugal Modulation in the Auditory System

Spectral processing is the fundamental function of auditory neurons throughout the central auditory system, which, in the ventral part of MGB and A1, is characterized by single peaked frequency tuning. Previous studies have shown that local inactivation of A1 altered the frequency tuning of MGB neurons by changing tuning bandwidth and/or shifting tuning peaks (Villa et al., 1991; Zhang et al., 1997). Consistent with these previous studies, we found that 48.5% of MGB neurons altered their frequency tuning when A1 was inactivated. Interestingly, during A1 inactivation, neurons with sharper tuning increased tuning bandwidth, whereas those with broader tuning decreased tuning bandwidth (Fig. 6G). These results suggest two roles for corticofugal modulation: one which increases frequency selectivity via sharpening of frequency tuning for increasing accuracy of sound perception, and another which broadens tuning bandwidth for greater spectral coverage, perhaps involving more complex integration.

### Working Model for Auditory Corticothalamic Modulation on MGB neurons

Based on the existing studies in auditory, visual, and somatosensory system of awake animals, we propose a model (Fig. 7) adapted from other sensory systems (Suga, 2020; Temereanca and Simons, 2004; Usrey and Sherman, 2019). In our model, neurons in A1 send both direct excitatory projections to MGB neurons as well as indirect extensive inhibitory projections to MGB neurons via thalamic reticular nuclear (TRN). Without auditory inputs, the balance of MGB neuronal response is dominated by global inhibition from TRN (path B); this serves to broadly suppress the background neural activity of MGB neurons, a suppression which is revealed by inactivation in A1 (Fig 2–3). During sound perception, A1 pushes the balance of activity towards greater excitation than inhibition (path A), thereby boosting the MGB response to the specific auditory input; this was revealed by our result that inactivation of A1 led to reduction of sound driven responses (Fig 2–3). The combination of these two corticofugal effects (similar to visual systems studies, excitation of ‘center’ response and suppression of ‘surround’ response) contribute to increase in SNR of MGB neurons during sound processing. In addition, consistent with previous experimental (Villa et al., 1991) and modelling studies (Suga, 2020), the balance between direct excitatory inputs and indirect inhibitory inputs determines whether the frequency tuning of the given MGB neuron is sharpened or broadened.

**Figure 7.**
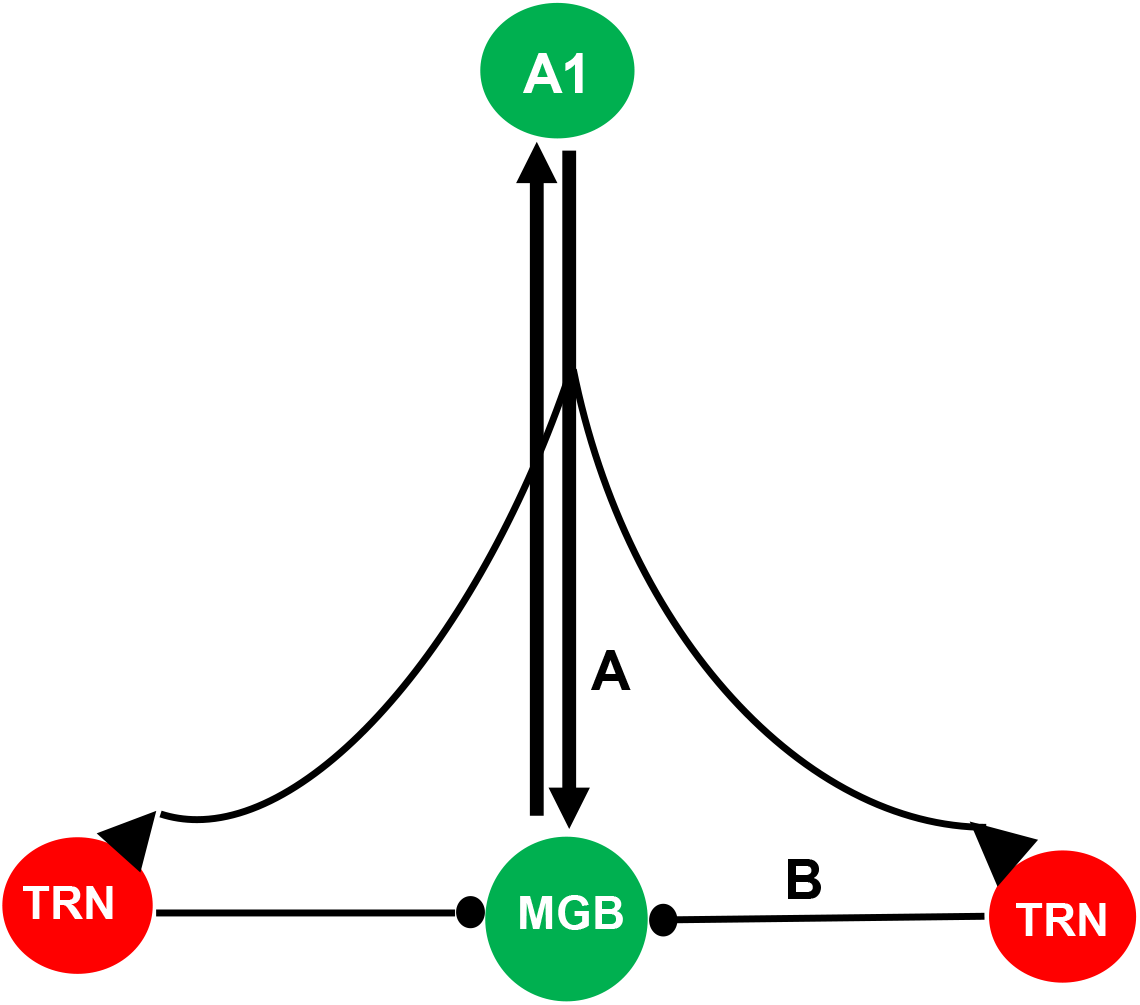
Proposed model of corticothalamic modulation. In our model, neurons in A1 send both direct excitatory projections to MGB neurons (path A) as well as indirect extensive inhibitory projections to MGB neurons via TRN (path B). Without auditory inputs, the balance of MGB neural response is dominated by global inhibition from TRN (path B); this serves to broadly suppress the background neural activity of MGB neurons. This was revealed by our result that inactivation of A1 led to an increase in spontaneous activity. During sound perception, A1 pushes the balance of activity towards greater excitation than inhibition (path A), thereby boosting the MGB response to the specific auditory input; this was revealed by our result that inactivation of A1 led to reduction of sound driven responses. The combination of these two corticofugal effects contribute to an increase in SNR of MGB neurons during sound processing. In addition, the balance between direct excitatory inputs and indirect inhibitory inputs determines whether the frequency tuning of the given MGB neuron is sharpened or broadened. A1, the primary auditory cortex; MGB, medial geniculate body; TRN, thalamic reticular nuclear.

Taken together, we find that the mammalian auditory system shares common corticofugal processing features with other sensory systems, something which in the auditory system is more apparent during information processing in the awake state.

## Acknowledgements

The authors would like to thank Anna Wang Roe for her critical reading of the early version of this paper. This work was supported by the Natural Science Foundation of China (61703365, 31871056, 91732302), National Key R&D Program of China (2018YFC1005003), and Fundamental Research Funds for the Central Universities (2018QN81008, 2019XZZX001-01-20).

## Author Contributions

L.G. designed the research; Y. Z adapted the cryo-inactivation method; Y. Z, X. W and L. Zhu collected the data; Y. Z and X. W. analyzed the data; X. W. plotted the figures; L.G. and X. L discussed the data and wrote the manuscript; Min Li helped with technical support for developing the cooling technique; others involved in discussion, editing, and helping with the technical support for electrophysiological recordings as well as animal preparations.

## Methods

All experimental procedures were approved by Animal Use and Care Committee of Zhejiang University following the National Institutes of Health (NIH) guidelines.

### Animal Preparation and Single-Unit Recording Procedures

Experiments were conducted in two female awake adult common marmosets (Callithrix jacchus, two females) using the chronic preparation (Lu et al., 2001). Detailed descriptions are in our previous study (Gao and Wang, 2020). In brief, the animals were trained to sit quietly in a custom-designed primate chair and training time was gradually increased from 15 min to 2 hours per day over the course of ~ 2 weeks until the animals sit quietly in the chair. A head-cap implantation surgery was performed under aseptic conditions, during which two headposts were attached on the skull of the animals in order to fasten its head for electrophysiological recordings. Two recording chambers were built with dental cement over the temporal side during the surgery and the lateral sulcus (LS) was traced as a landmark to help identify the location of the auditory cortex. To access A1 and MGB, small craniotomies (1mm in diameter) were made in the recording chambers to allow for the penetration of electrodes. The A1 was identified by its tonotopy map along the LS at an angle of 60° (Bendor and Wang, 2008). The recording electrode approached the MGB in a dorsolateral-to-ventromedial trajectory, entering the brain 3 mm lateral to the high-frequency representation of A1 along the LS at an angle of 57.5°. The auditory thalamic neurons were encountered ~ 6–9 mm beneath the cortical surface (Bartlett and Wang, 2011). Single-unit recordings were made using high impedance tungsten microelectrodes (2–5 MΩ, FHC) advanced by a one-axis motorized stereotaxic micromanipulator (DMA1510, NARISHIGE). The signals were amplified (AlphaLab SNR, Alpha Omega Engineering) and digitized (RX6, Tucker-Davis Technologies), saved and analyzed using custom programs written in MATLAB (Mathworks). The spikes were detected on-line using a template-matching method (AlphaLab SNR, Alpha Omega Engineering). The animal was head-fixed and semi-restrained in a custom-made marmoset chair, but was not performing a task during these experiments.

Male and female Sprague Dawley (SD) rats (250–350 g) with clean external ears were used to validate the cooling effect of the cryoloop. The animals were anesthetized with urethane (1.2 g/kg, 20%) via intraperitoneal injection and maintained with supplementary doses (0.2 g/kg/h) as needed. The level of anesthesia was monitored frequently by paw pinch and corneal reflexes. Atropine sulfate (0.05 mg/kg) was administered subcutaneously 15 minutes prior to the anesthesia to suppress tracheal secretion. 2% of lidocaine was applied locally under the skin before a midline incision. A headpost was attached to exposed skull with dental cement for head fixation during single unit recordings. The animal was then placed inside a soundproof chamber, and its head was fixed by the headpost. The body temperature of the animals was maintained by a feedback-controlled heating pad. To access the auditory cortex, one craniotomy (5 mm x 1.5 mm) was made on the left temporal side that overlays the primary auditory cortex (Gao et al., 2017).

### Acoustic Stimuli

All recording sessions were carried out in a double-walled, soundproof chamber with inside dimensions of 2.8 m X 2.8 m X 1.9 m, which has a 50 dB reduction of environment noise on average (FOSHANHENQI). Acoustic signals were generated digitally at a sampling rate of 97.7 kHz using custom MATLAB software (MathWorks), low-pass filtered at 48.8 kHz, converted to analog signals (RX6, Tucker-Davies Technologies), power amplified (PM5005, Marantz), attenuated with two serially linked attenuators (PA5, Tucker-Davies Technologies), and delivered in free-field through a speaker (8351A, Genelec) located approximately 1.2 m in front of the animal’s head. The measured speaker output was within 6 dB of the intended output for frequencies from 32 Hz to 40 kHz, which encompasses the hearing range of marmoset monkeys, with a calibrated sound level of 90 dB sound pressure level (SPL) at 0 dB attenuation for 1-kHz tones.

Once a neuron in MGB/A1 was isolated, pure tone and white broad-band Gaussian noise were played to characterize its basic response properties, such as best frequency (BF), best level (BL). Broad-band Gaussian noise (FIR filtered, flat spectrum between 0.25 - 40 kHz, −10 to 70 dB SPL) with varying sound level were played to characterize neuron’s rate level function and preference to tone or noise. Pure tones (frequency: 0.25-40 kHz, duration: 200 ms, 5 ms cosine ramps) or narrow-band Gaussian noise (center frequency: 0.25-40 kHz, duration: 200 ms, bandwidth: 1 octave) were presented spanning 3–4 octaves around a manually determined center frequency in 0.1-octave in randomized blocks. The range of the sound level was from −10 to 70 dB SPL. Sinusoidal amplitude modulated tone or noise were played as a time-varying stimulus in randomized blocks, in which the carrier frequency set at a neuron’s BF, was held constant while its amplitude was modulated by a sinusoid signal and modulation frequency (MF) varied between 2 and 512 Hz on a logarithmic scale.

### Cryoloop Cooling Apparatus

The cooling method for awake marmosets was adopted from that developed in anesthetized cats by Lomber et.al (Lomber et al., 1999), and is a modified and miniaturized version. The cryoloop is manufactured from 17-gauge (1.4 mm O.D. x 1.1 mm I.D.) hypodermic stainless-steel tubing, which was then shaped into a loop to allow it to conform to the surface of marmosets A1. A microthermocouple (TC), made by twisting Teflon insulated copper (38 AWG gauge, 0.102 mm) and constantan wire (Omega Engineering Limited) together and trimming the tips closely, is constructed and attached to the cryoloop. The wire of TC was connected to a DC power temperature measuring system to monitor the change of temperature on the cortical surface in real time. The TC has both high resolution and fast response time in the operating range of 0 – 40 °C. The inlet and outlet of the cryoloop connected with silicone tubing, 1 m of which was buried into dry ice before reaching the cryoloop. The ends of silicone tubing came into a reservoir filled with ethanol (histological grade), which was driven by a peristaltic pump (Longer Pump, #13). When the cooling system was turned on, the chilled ethanol passed through the cryoloop. The temperature of brain tissue underneath the cryoloop was controlled by velocity and duration of chilled ethanol flow passing through the cryoloop tubing system as shown in Figure 1C. The Cryoloop assembly was sterilized with ethylene oxide before use. We examined the cooling effects by single unit recordings after the cryoloop was positioned in A1 by a stereotaxic micromanipulator (SM11, NARISHIGE) via a craniotomy made over A1. The tips of the TC were placed on the surface of A1. Next, the cryoloop assembly was fastened into the recording chamber by dental cement (UNIFAST, Trad, GC Corporation) after filling the craniotomy with silastic (KWIK-SIL, World Precision Instruments). Notably, the cryoloop must be fitted closely to the brain surface to ensure the validity of inactivation effect.

### Data Analysis

The analysis methods were basically the same as in previous publications (Bendor and Wang, 2008; Gao and Wang, 2019). Spontaneous rate was calculated as the mean firing rate before sound presentation over the entire stimulus set unless otherwise specified; driven rates were calculated over the entire stimulus duration and the spontaneous rate was subtracted in all analyses unless otherwise specified. The onset response was defined as driven rate 0 - 40 ms from sound stimulation onset whereas the sustained response was defined as driven rate 40 - 200 ms from sound stimulation onset. The criterion for a significant stimulus-driven response was defined as averaged driven rate that was larger than 2 standard deviations (2SDs) above the mean spontaneous firing rate. BF was defined as the tone frequency that generated the highest firing rate in the recorded neuron. Non-BF was defined as the frequency which elicited the smallest statistically significant driven response. The tuning bandwidth (in octaves) of MGB neurons was taken as the width at half maximum firing rate after we plotted the frequency tuning curve at best level or 40 dB SPL in 0.1-octave steps. Best modulation frequency (rBMF) was defined as the modulation frequency of sAM that generated the highest firing rate in the recorded neuron. Response latency was calculated by finding the earliest time after stimulus onset where three consecutive 2-ms bins had discharge rates 2SDs above the mean spontaneous discharge rate. The signal-to-noise ratio was defined as 20* log_10_ (averaged driven rate/ averaged spontaneous rate). If a neuron has no spontaneous activity, the maximum value of SNR among all the neurons in a certain condition (before or during A1 inactivation) was set as the SNR of this neuron. If the driven rate of a neuron is zero, the minimal value of SNR among all the neurons in the same condition (before or during A1 inactivation) was set as SNR of this neuron.

Change (decreased or increased) of SNR was calculated by the formula as follows: F = (C – B) /B, where C is the SNR during A1 inactivation; B is the SNR before A1 inactivation. Increased SNR is defined as F>0.05; decreased SNR is defined as F<0.05; otherwise, it is defined as no change. Change (decreased or increased) of tuning bandwidth or BF was defined as those outside the confidence interval of ± 0.2 octaves. Change index (F) was used to quantify the degree of change in sound driven rate and spontaneous firing rate of MGB neurons caused by inactivation of A1. The formula is as follows: F = (C − B) / (C + B), where C is the sound driven rate of MGB neuron arising from certain acoustic stimulation during inactivation of A1; B is the corresponding value before inactivation of A1. If F <−1, we set it as −1. If F >1, we set it as 1. Significant change or difference was defined as being greater than 0.05 unless otherwise stated.

All values are expressed as mean ± sem (standard error of the mean) unless otherwise specified. Student’s paired or unpaired t test was used for an analysis of the data. P values with p<0.0001, p<0.001, p<0.01, or p<0.05 were considered statistically significant in respective tests, as indicated where appropriate.

## Supplemental Information

Supplemental Information includes 2 Supplemental figures

**Figure S1. Validation of Effectiveness of Cooling Method in Auditory Cortex of Anesthetized Rats.**

(**A**) Raster plots showing the neural responses to white broad-band noise (WB) with varying intensities in the auditory cortex of anesthetized rats when the auditory cortex was cooled to different temperatures.

(**B**) Mean driven rate of auditory cortical neurons in response to WB at 70 dB SPL plotted against temperature detected at the surface of the auditory cortex (32 °C, n=11; 27 °C, n=9; 22 °C, n=10; 17 °C, n=11; 13 °C, n=11; 8 °C, n=11; 3 °C, n=11; 32 °C, n=8).

(**C-E**) Raster plots showing neural responses to WB for a sampled cortical neuron at different depth (C, 0.5 mm; D, 2 mm; E, 2.5 mm) from the auditory cortical surface of anesthetized rats before, during and after the auditory cortex was cooled to 3 °C.

(**F-H**) Raster plots showing neural responses to WB for three sampled neurons with different distances from the edge of the cryoloop (distance: F, 0.5 mm; G, 2 mm; H, 2.5 mm) before, during and after the auditory cortex was cooled to 3 °C. Green shaded areas in A-H indicate time periods of acoustic stimulation.

(**I**) Mean driven rate to WB at 70 dB SPL were plotted against the recording depths. For neurons without auditory response, firing rate in entire recording duration (0-700 ms) was calculated and used in qualification (0.5mm, n=6; 1mm, n=9; 1.5mm, n=5; 2 mm, n=2; 2.5mm, n=2).

(**J**) Mean driven rate to WB at 70 dB SPL were plotted against recording distance from the edge of the cryoloop (0.5 mm, n=6; 1 mm, n=7; 1.5 mm, n=8; 2 mm, n=6; 2.5 mm, n=4).

**Figure S2. Auditory Response of Some MGB Neurons Disappeared When A1 was Inactivated**

Raster plots of neural response for four example MGB neurons in response to pure tones with varying frequencies before (Left), during (Middle) and after (Right) inactivation of A1. Green shaded areas indicate time periods of acoustic stimulation.

