## Supplementary figures and images for "Selective Corticofugal Modulation on Sound Processing in Auditory Thalamus of Awake Marmosets"

### Supplemental figure 1

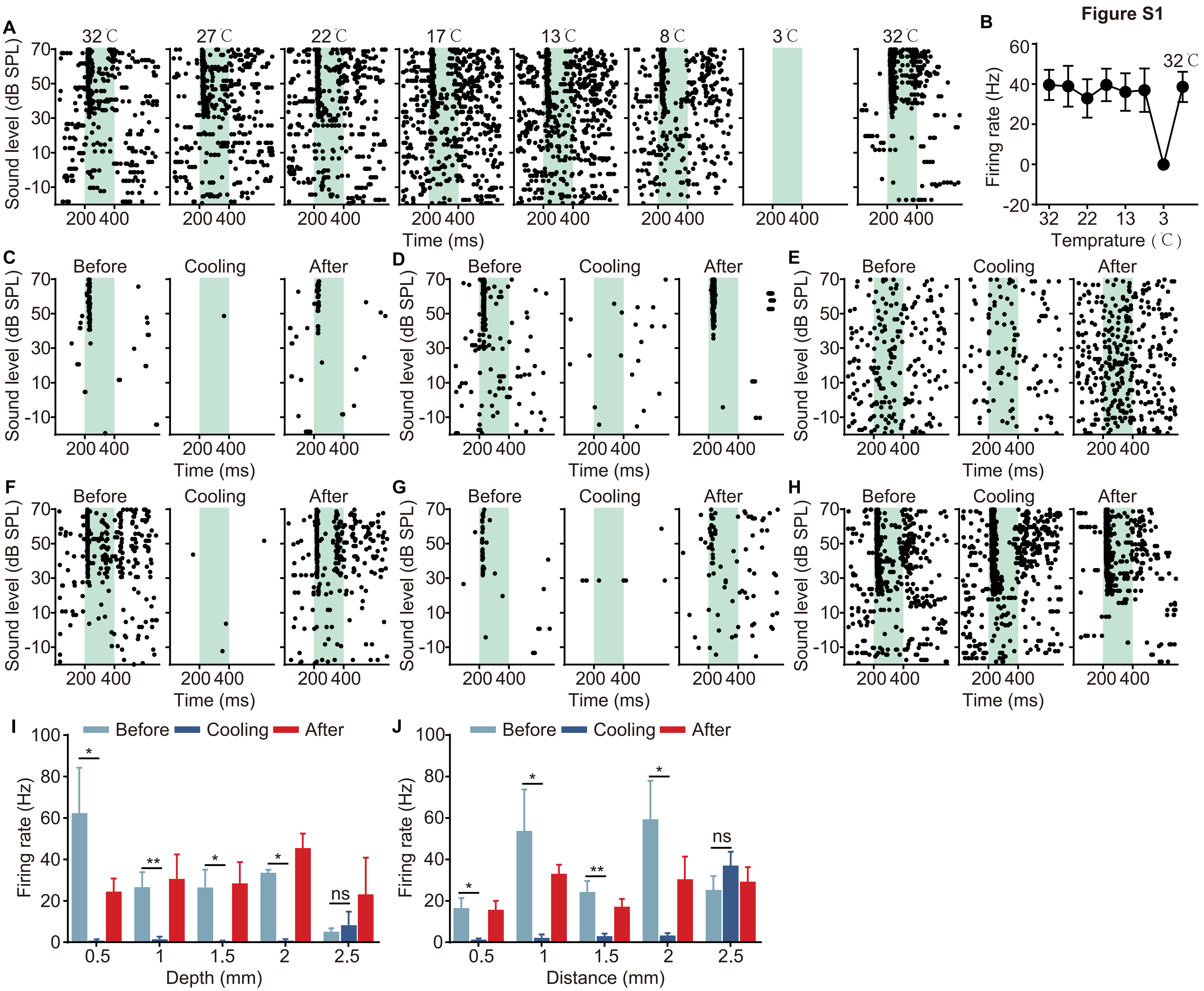

### Supplemental figure 2

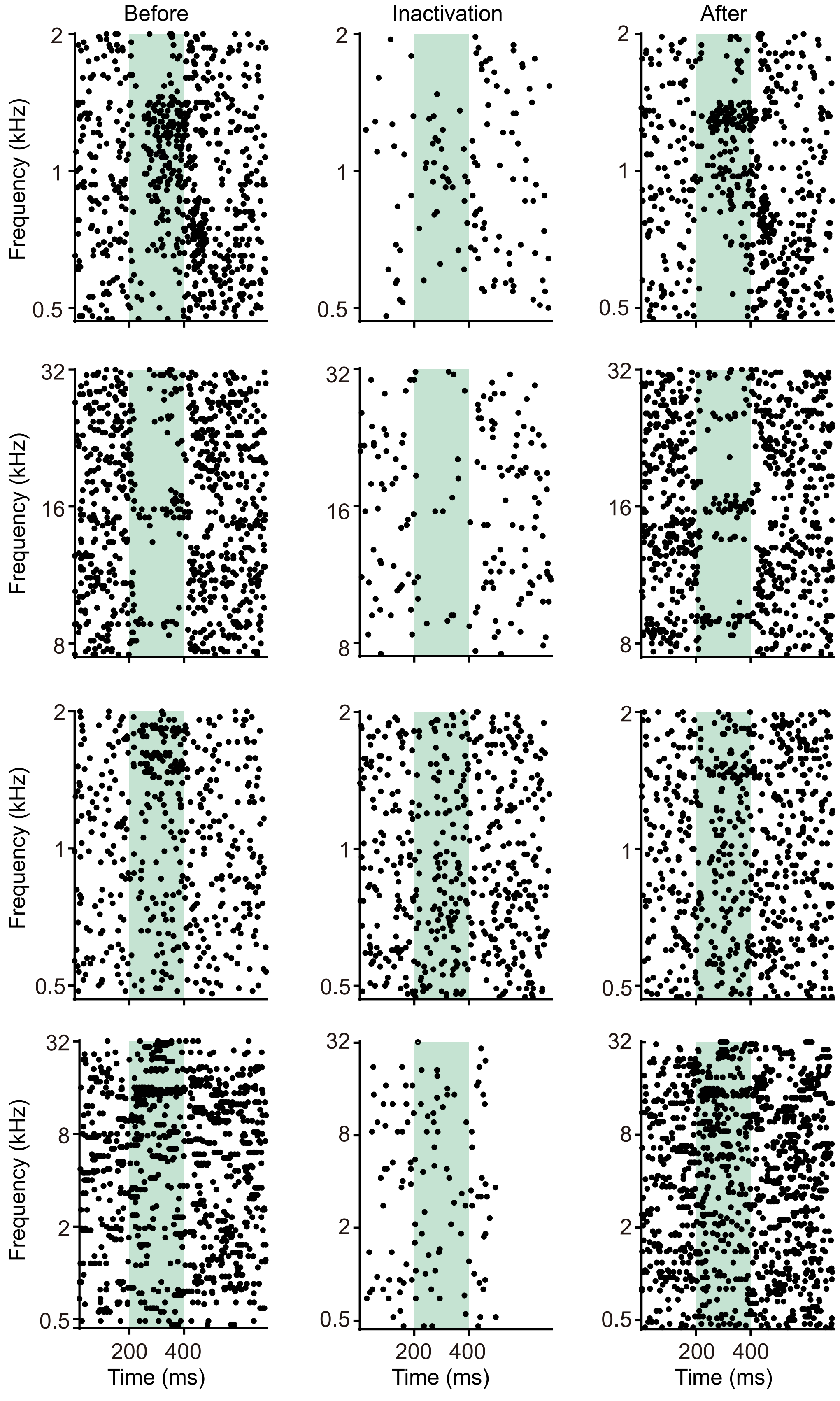
